# Perception of dynamic facial expressions activates a cortico-subcortico-cerebellar network in marmosets

**DOI:** 10.1101/2022.08.02.502528

**Authors:** Audrey Dureux, Alessandro Zanini, Stefan Everling

**Author notes:** <u>Corresponding author:</u> Audrey Dureux, Centre for Functional and Metabolic Mapping, Robarts Research Institute, University of Western Ontario, London, Canada.

## Abstract

The perception and identification of faces and facial emotional expressions are critical for social communication in the daily life of all primates. Here, we investigated the neural network activated by dynamic facial expressions in awake New World common marmosets with ultra-high field fMRI at 9.4T. Our results show that dynamic facial expressions activate several areas along the occipitotemporal axis (V2/V3, V4/TEO, FST, caudal TE, rostral TE, TPO), frontal cortex (areas 45/47, 13, 8a and orbital cortex), amygdala, motion-sensitive areas, premotor area, and in the posterior lobe of the cerebellum. Negative faces increased the monkeys’ respiration rates and elicited stronger responses along the occipitotemporal cortical axis, in the amygdala and in the hypothalamus. This cortico-subcortico-cerebellar network may play an important role in the perception of behaviorally relevant facial expressions that are vital for social communication in marmosets.

**Significance Statement:** Recent research has highlighted the importance of emotional content of faces in social communication in humans and non-human primates. The current study focuses on the neural responses to dynamic emotional facial expressions in the common marmoset (Callithrix jacchus), a New World primate species sharing several similarities of social behavior with humans. Using ultra-high-field fMRI, we show that negative facial expressions activate a cortico-subcortico-cerebellar network and, critically, negative faces increase the level of arousal of marmosets, possibly relayed through a modulation of the activity in the autonomic nervous system via stress-integrative brain centres in the hypothalamus.

Our results reveal the existence of specific neural and physiological responses to negative emotional faces suggesting that behaviorally relevant facial expressions are vital for social communication in New World marmosets.

## Introduction

Social cognition requires the ability to perceive, process and recognize the emotional expressions of conspecifics (Ferretti and Papaleo, 2019). This ability is not only crucial for humans but also for a number of other social species (Albuquerque et al., 2016; Parr et al., 1998; Tate et al., 2006) allowing them to communicate intentions and to better adapt to environmental challenges (Russell, 1997). Facial expressions are particularly necessary during social interactions, constituting a powerful way for rapid nonverbal communication in all primates (Darwin, 2004).

Functional magnetic resonance imaging (fMRI) studies have identified the neural networks underlying face and facial expression processing in humans and non-human primates. In humans, faces elicit activity in a cortical and subcortical network that includes the fusiform gyrus, amygdala and fronto-temporal regions in anterior and posterior regions of the superior temporal sulcus (STS) and in the inferotemporal cortex as well as orbitofrontal cortex (Fusar-Poli et al., 2009; Haxby et al., 2000; Kanwisher et al., 1997; Puce et al., 1998). Distinct patterns of activation have also been associated with different facial expressions. Particularly, the recognition of fearful faces activates amygdala, visual cortex (Blair, 2003; Vuilleumier and Pourtois, 2007) for a review), and the lateral orbitofrontal cortex (Murphy et al., 2003), whereas the observation of angry faces activates the insula, cingulate, thalamus, basal ganglia, and hippocampus (Strauss et al., 2005). In general, the processing of emotional faces compared to neutral faces is associated with increased activation in a set of visual, temporoparietal, limbic and prefrontal areas as well as the putamen and the cerebellum (Fusar-Poli et al., 2009). These brain areas respond even more when dynamic facial expressions are used compared to static ones, which also seem to facilitate their recognition (Ceccarini and Caudek, 2013; Zinchenko et al., 2018).

Comparable face-selective regions have been identified in Old World macaque monkeys (Tsao et al., 2003) and in New World marmosets (Hung et al., 2015; Schaeffer et al., 2020). In macaques, face-selective patches are located along the occipitotemporal axis in occipital cortex, along the STS, in ventral and medial temporal lobe, in some subregions of the inferior temporal cortex, as well as in frontal cortex and some subcortical regions such as amygdala and hippocampus (Bell et al., 2009; Hadj-Bouziane et al., 2008; Pinsk et al., 2005; Tsao et al., 2003; Weiner and Grill-Spector, 2015). In marmosets, face patches are found along the occipitotemporal axis and in frontal cortex (Hung et al., 2015; Schaeffer et al., 2020). Some fMRI studies have also revealed a stronger modulation of face-selective regions and surrounding cortex for emotional compared to neutral faces in Old World macaques (Hadj-Bouziane et al., 2008; Hoffman et al., 2007; Liu et al., 2017). Furthermore, lesion studies in both humans and macaques suggest that the amygdala plays a key role in the recognition of facial expressions (Adolphs et al., 1994; Aggleton and Passingham, 1981; Hadj-Bouziane et al., 2012).

Here, we investigated the neural circuitry involved in the processing of dynamic facial expressions in marmosets using ultra-high-field (9.4T) fMRI. We acquired whole brain fMRI in six awake marmosets while they viewed videos of marmoset faces with neutral or negative facial expressions. To further explore the physiological state of the monkeys during the observation of these different emotional contexts, we also recorded the animals’ respiratory rate inside the scanner during the task.

## Materials and methods

### Common Marmoset Subjects

For this study we acquired ultra-high field fMRI data from six awake common marmoset monkeys (*Callithrix jacchus*): three females (MGR, MHE, MKO: weight 345-442 g, age 30-34 months) and three males (MMI, MMA, MGU: weight 348-429 g, age 30-55 months). All experimental procedures follow the guidelines of the Canadian Council of Animal Care policy and a protocol approved by the Animal Care Committee of the University of Western Ontario Council on Animal Care #2021-111.

### Surgical procedure

To prevent head motion during MRI acquisition, the marmosets were surgically implanted with an MR-compatible head restraint/recording chamber (MGR, MHE, MMI, MMA and MGU) or with an MR-compatible head post (MKO), conducted under anesthesia and aseptic conditions.

During the surgical procedure, the animals were firstly sedated and intubated to maintain the animals under gas anaesthesia with a mixture of O_2_ and air (isoflurane 0.5-3%). With their head immobilized in a stereotactic apparatus, the head fixation device was positioned on the skull after a midline incision of the skin along the skull and maintain in place using a resin composite (Core-Flo DC Lite; Bisco). Additional resin was added as needed to cover the skull surface and ensure an adequate seal around the device. For optimal adhesion, the skull surface was well prepared before the application of the resin by applying two coats of an adhesive resin (All-Bond Universal; Bisco, Schaumburg, IL) using a microbrush, air-dried, and cured with an ultraviolet dental curing light (King Dental) (for more details, see (Johnston et al., 2018)). Throughout the surgery, heart rate, oxygen saturation, and body temperature were continuously monitored. Two weeks after the surgery, the monkeys were acclimatized to the head-fixation system and the MRI environment in a mock scanner.

### Experimental setup

During the scanning sessions, monkeys sat in a sphinx position in the MRI-compatible restraint system (Schaeffer et al., 2019) positioned within a horizontal magnet (9.4 T MR scanner). Their head was restrained by the fixation of the head chamber or the head post. An MR compatible camera (model 12M-i, 60 Hz sampling rate, MRC Systems GmbH, Heidelberg, Germany) was positioned in front of the animal allowing to monitor the animal during the acquisition.

Inside the scanner, monkeys faced a plastic screen placed at 119 cm from the animal’s head where visual stimuli were projected with an LCSD-projector (Model VLP-FE40, Sony Corporation, Tokyo, Japan) via a back-reflection on a first surface mirror. Visual stimuli were presented with Keynote software (version 12.0, Apple Incorporated, CA) and were synchronized with MRI TTL pulses triggered by a Raspberry Pi (model 3B+, Raspberry Pi Foundation, Cambridge, UK) running via a custom-written python program. No reward was provided to the monkeys during the scanning sessions.

### Stimuli

Visual stimuli consisted of video clips showing faces of marmosets depicting either neutral or negative facial expressions as well as scrambled version of each. We recorded these videos from six marmosets while they sat non-head-fixed in a marmoset chair (Johnston et al., 2018). To obtain the negative facial expression, a plastic snake was presented to the monkeys while they were filmed. Then, 12-sec video clips were created using custom video-editing software (iMovie, Apple Incorporated, CA). Scrambled versions of these videos were created by random rotation of the phase information using a custom program (Matlab R2022a, the Mathworks). The same random rotation matrix was used for each frame in the scrambled conditions to preserve motion and luminance components.

### Passive viewing facial emotional expressions task

A block design was used in which each run consisted to eight blocks of stimuli (12 sec each) interleaved by nine baseline blocks (18 sec each; see Figure 1). In these blocks, four different conditions were repeated twice. These conditions consisted to video clips of marmoset’s face: 1) neutral faces, 2) negative faces, 3) scrambled neutral faces, and 4) scrambled negative faces. The order of these conditions was randomized between each run leading to 20 different stimulus sets, counterbalanced within the same animal and between animals. During baseline blocks, a 0.36° circular black cue was displayed in the center of the screen against a gray background. We found previously that such a stimulus reduced the vestibulo-ocular reflex evoked by the ultra-high field MRI. To compare the physiological state of each animal during the different conditions, we also measured respiration rate (RR) during the runs via a sensor strap (customized respiration belt) around the chest of the animals and recorded the signals with the LabChart software (ADInstruments).

**Figure 1.**
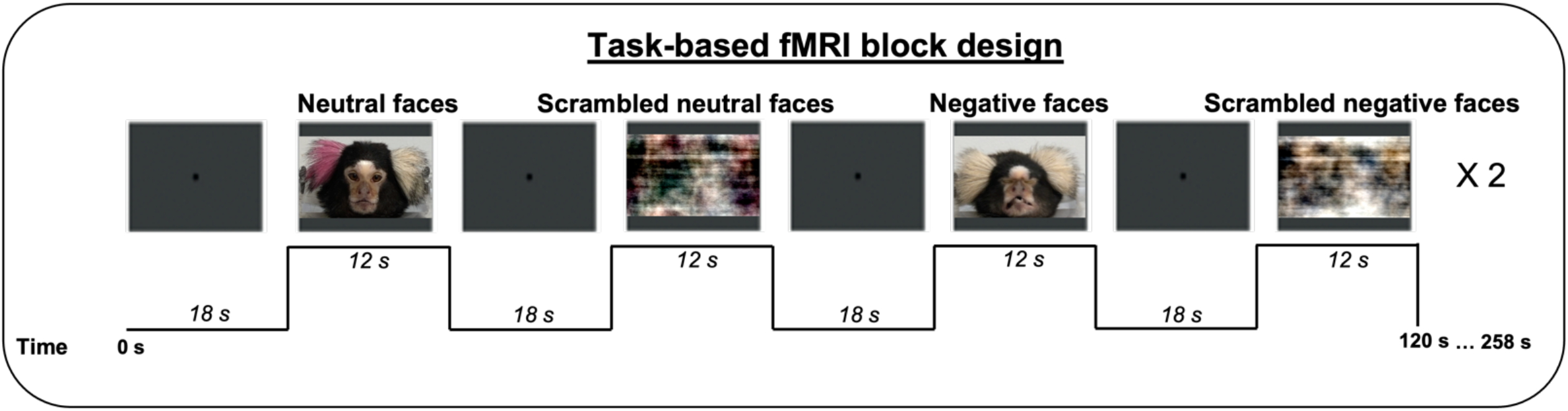
Passive viewing fMRI task block design. In each run, four different types of videos lasting 12 sec each were presented twice in a randomized order. The videos consisted of faces of marmosets depicting different facial expressions (neutral faces and negative faces conditions) and their scrambled version (scrambled neutral faces and scrambled negative faces conditions). Each video was separated by baseline blocks of 18 sec, where a central dot was displayed in the center of the screen.

### fMRI data acquisition

Data were acquired on a 9.4 T 31 cm horizontal bore magnet and Bruker BioSpec Avance III console with software package Paravision-6 (Bruker BioSpin Corp). We used a custom-made five channel (subjects MGR, MHE, MMI, MMA and MGU) or eight channel (subject MKO) receive surface coil paired with a custom-built high-performance 15-cm-diameter gradient coil with 400 mT/m maximum gradient strength (xMR, London, Canada; Peterson et al., 2018). For the radiofrequency transmit coil, we used a quadrature birdcage coil (12 cm inner diameter) built and customized in-house.

Functional gradient-echo-based single-shot echo-planar images (EPI) covering the whole brain were acquired over multiple sessions including a total of 79 runs for all 6 animals (9-19 runs/animal depending on the compliance of the animal), using the following sequence parameters: TR=1.5s, TE = 15ms, flip angle = 40°, field of view=64×48 mm, matrix size = 96×128, resolution of 0.5 mm3 isotropic, number of slices= 42 [axial], bandwidth=400 kHz, GRAPPA acceleration factor: 2 (left-right). Another set of EPI with an opposite phase-encoding direction (right-left) has been collected for the EPI-distortion correction. For each animal, a T2-weighted structural was acquired during one of the sessions (TR=7s, TE=52ms, field of view=51.2×51.2 mm, resolution of 0.133×0.133×0.5 mm, number of slices= 45 [axial], bandwidth=50 kHz, GRAPPA acceleration factor: 2.

### fMRI data preprocessing

fMRI data were preprocessed with AFNI (Cox, 1996) and FMRIB/FSL (Smith et al., 2004) software packages. After converting the raw functional images into NifTI format using dcm2nixx function of AFNI, the images were reoriented from the sphinx position using FSL (FSL’s fslswapdim and fslorient functions). Functional images were then despiked (AFNI’s 3Ddespike function) and volume were registered to the middle volume of each time series (AFNI’s 3dvolreg function). The motion parameters from volume registration were stored for later use with nuisance regression. Images were then smoothed at full width at half-maximum Gaussian kernel (FWHM) of 2mm (AFNI’s 3dmerge function) and bandpass filtered (0.1 to 0.01 Hz) to prepare the data for the regression analysis (AFNI’s 3dBandpass function). An average functional image was calculated for each run and linearly registered to the respective T2-weighted anatomical image of each animal (FSL’s FLIRT function). The transformation matrix obtained after the registration was then used to transform the 4D time series data. T2-weighted anatomical images were manually skull-stripped, and the mask of each animal was applied to the corresponding functional images.

Finally, T2-weighted anatomical images were registered to the NIH marmoset brain atlas (Liu et al., 2018) via the nonlinear registration using Advanced Normalization Tools (ANTs’ ApplyTransforms function).

### Statistical analysis of fMRI data

For each run, the task timing was convolved to the hemodynamic response (AFNI’s ‘BLOCK’ convolution) and a regressor was generated for each condition to be used in a regression analysis (AFNI’s 3dDeconvolve function). All four conditions were entered into the same model, along with polynomial detrending regressors and the motions parameters.

The resultant regression coefficient maps were then registered to template space using the transformation matrices obtained with the registration of anatomical images on the template (see above). We therefore obtained four T-value maps for each run registered to the NIH marmoset brain atlas (Liu et al., 2018). These maps were then compared at the group level via paired t-tests using AFNI’s 3dttest++ function, resulting to Z-value maps. To protect against false positives, a clustering method derived from 10000 Monte Carlo simulations was applied to the resultant t-test maps using ClustSim option (*α*=0.05).

To identify brain regions implicated in facial expression processing in general, we first examined the voxels that were significantly stronger activated by the combination of all facial expressions conditions compared to the combination of all scrambled facial expressions conditions (i.e., neutral + negative faces > scrambled neutral + scrambled negative faces contrast). Then, to investigate voxels dedicated to the two types of facial expressions (negative and neutral), we compared the activations obtained for each facial expression with their scrambled version (i.e., neutral faces > scrambled neutral faces and negative faces > negative scrambled faces contrasts). Finally, we identified voxels more activated by negative facial expressions compared to neutral facial expressions (i.e., negative faces > neutral faces).

The resultant Z-value maps were displayed on fiducial maps obtained from the Connectome Workbench (v1.5.0, Marcus et al., 2011) using the NIH marmoset brain template (Liu et al., 2018) and on coronal sections. We used the Paxinos parcellation of the NIH marmoset brain atlas (Liu et al., 2018) to define anatomical locations of cortical and subcortical regions.

Based on the existing fMRI literature on faces processing in marmosets (Hung et al., 2015; Schaeffer et al., 2020) and facial expression processing in humans and macaques (Fusar-Poli et al., 2009; Hadj-Bouziane et al., 2012, 2008; Vuilleumier and Pourtois, 2007), fourteen regions of interest (ROIs) were also selected from the Paxinos parcellation of the NIH atlas (Liu et al., 2018): occipitotemporal areas V2, V3, V4, V4T, TEO, FST, TE1, TE2, TE3; frontal areas 45, 47M, 47O, 47L and amygdala as subcortical areas). First, bilateral masks of these ROIs were performed using AFNI’s 3dcalc function using the Paxinos parcellation of the NIH atlas (Liu et al., 2018). Second, times series for each condition and each run were extracted from the resultant regression coefficient maps using AFNI’s 3dmaskave function. Finally, the difference in response magnitude (i.e., percentage of signal change) was obtained for each condition and the difference in magnitude of these responses between conditions was computed using one sided paired t-tests with FDR post-hoc correction (p < .05) in custom-written Matlab scripts (R022a, The Mathworks).

### Respiratory rate analyses

A customized respiratory belt around both the ventral and dorsal thorax of the marmoset was used to record respiratory signals. The strength of these signals could vary between the different sessions as the positioning of the belt could change with the animal’s movement. Therefore, we preliminarily inspected the quality of the data and excluded all runs where segments of signals didn’t demonstrate stable respiratory signals.

For each animal, we decided to only include the first run of each session in order to avoid any habituation effect which may cause a decreased response of respiratory rate induced by the repeated viewing of the negative facial expression across a same session. Indeed, it has been shown that fear responses, particularly the physiological responses, decline over repeated exposures to fear-provoking stimuli (van Hout and Emmelkamp, 2002). Therefore, a total of 17 runs were selected throughout all the sessions for the 6 animals (with a minimum of 2 runs per animal).

The respiration signals recorded during the experiments were analyzed with LabChart data analysis software (ADInstruments) to obtain average respiratory rate (RR) per minute (bpm) for each animal in each block of the runs. For that, we first detected all the inter-beat RR peaks automatically. The number of R peaks for 1 minute correspond to the number of respirations so the number of times the chest rises (beats) in one minute (bpm), i.e., RR frequency. Then, we computed the average number of breaths per minute (i.e., mean RR) during the period of block of stimulus presentations (12 sec of video clips) and also during the baseline periods (central cross block of 18 sec).

External triggers were recorded for every TR (i.e., every 1.5 sec). The first trigger started as soon as the central dot appeared, and we extracted RR data for the baseline period counting 12 triggers (corresponding to 18 sec). Then, we extracted RR data for the video clip periods during the 8 following triggers (corresponding to 12 sec) and we repeated the same steps until the end of the run. In this way we obtained RR values for 9 blocks of baseline and 8 blocks of video clips for each run. We then computed the mean RR for each condition in each run (i.e., neutral faces, negative faces, scrambled neutral faces and scrambled negative faces) as well as the mean RR for the entire period of baseline for each run. Finally, we computed the mean delta RR by subtracting each mean RR obtained for each condition by mean RR obtained during the baseline period. As the respiratory rate range was variable between animals, we computed the delta RR in order to normalize these ranges between animals. Then, we were able to determine the effect of dynamic facial expression on animals’ RR and to compare these fluctuations between conditions for the group.

To identify the effect of facial expression on respiratory signals, three paired t-tests were used on delta RR (p < .05): neutral faces *vs*. scrambled neutral faces, negative faces *vs*. scrambled negative faces and negative *vs*. neutral. To investigate potential habituation effects by negative faces in physiological signals, we also performed the same analysis by selecting only last runs of each session (total of 17 runs throughout all the sessions for the 6 animal). Four paired t-tests were performed on delta RR to compare each condition between first and last runs: first runs neutral faces *vs*. last runs neutral faces, first runs negative faces *vs*. last runs negative faces, first runs scrambled neutral faces *vs*. last runs scrambled neutral faces and first runs scrambled negative faces *vs*. last run scrambled neutral faces. Data were analyzed with the open-source software R (The R Core Team, 2013).

## Results

We presented four different conditions in a block-design fMRI task in six awake marmosets (i.e. (1) neutral faces, (2) negative faces, (3) scrambled neutral faces, and (4) scrambled negative faces) to characterize the brain network involved in dynamic emotional faces processing and to identify brain areas dedicated to the processing of negative facial expression. We also measured the monkeys’ physiological state to identify the effect of negative facial expression on their arousal level during the fMRI scans.

### fMRI results

#### Dynamic emotional faces recruit a cortico-subcortico-cerebellar brain network

First, we identified the brain regions that are more active during the observation of dynamic emotional faces compared to the scrambled emotional faces (i.e., all emotional faces > all scrambled faces contrast). Figure 2A shows group activation maps obtained by comparing the emotional faces video conditions (including both neutral and negative faces) and scrambled versions of those videos.

**Figure 2.**
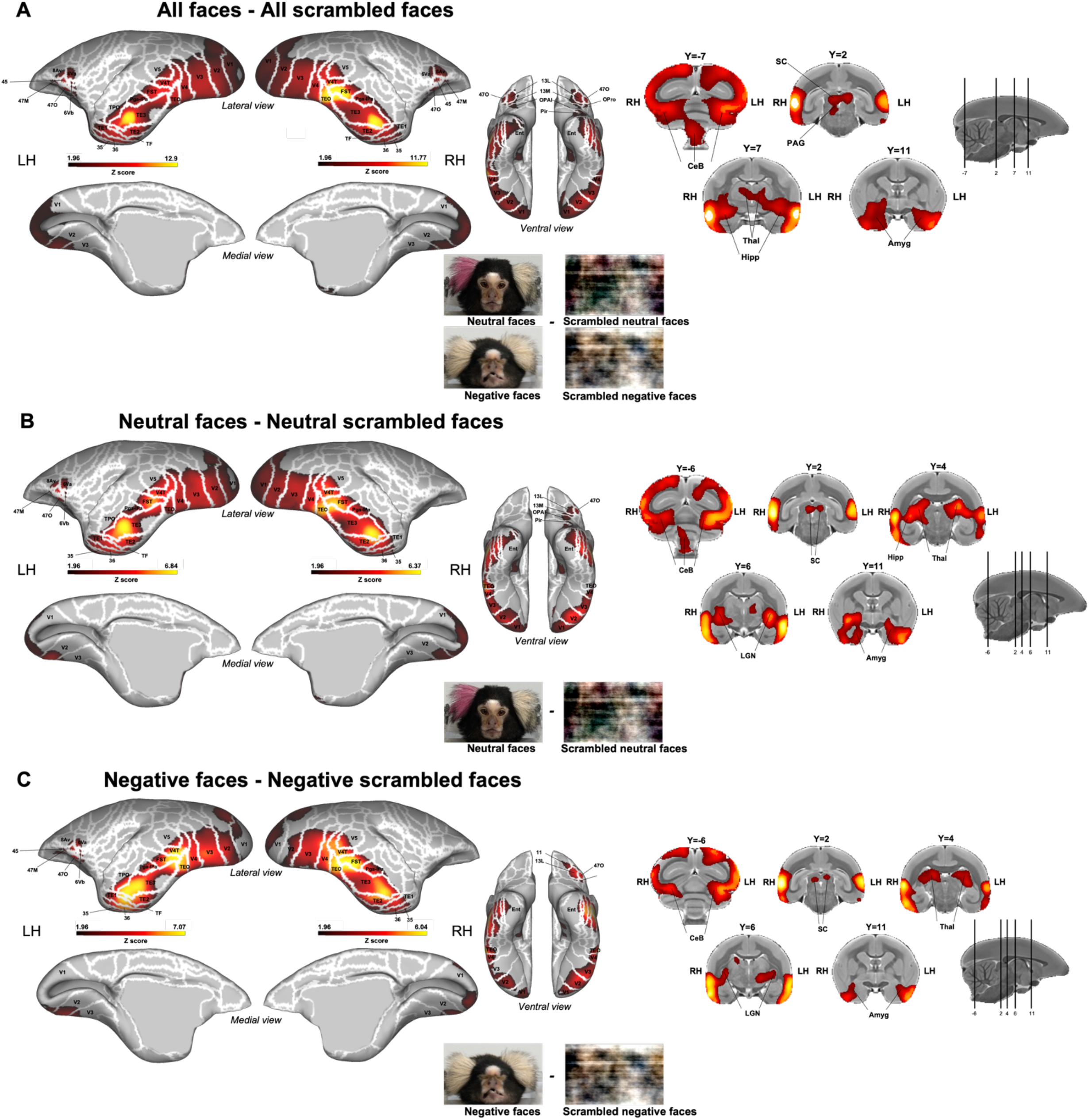
Brain networks involved in dynamic facial expressions processing. **A**. Group functional maps showing significant greater activations for the comparison between all emotional faces (i.e., neutral and negative facial expressions) and all scrambled emotional faces (i.e., scrambled versions of neutral and negative facial expressions). **B**. Group functional maps showing significant higher activations for neutral facial expression as compared to scrambled version of neutral faces. **C**. Group functional maps depicting brain regions with significant greater activations for the comparison between negative facial expression and scrambled version of negative facial expression. Group functional topology comparisons are displayed on the left and right fiducial marmoset cortical surfaces (lateral, medial and ventral views) as well as on coronal slices, to illustrate the activations in subcortical areas. The white line delineates the regions based on the Paxinos parcellation of the NIH marmoset brain atlas (Liu et al., 2018). The brain areas reported have activation threshold corresponding to z-scores > 1.96 (p<0.05, AFNI’s 3dttest++, cluster-size correction *α*=0.05 from 10000 Monte-Carlo simulations).

Significant preferential activations are observed for all faces compared to all scrambled faces in a large network comprising a set of bilateral cortical and subcortical regions. The major significant clusters are found along the occipitotemporal axis in visual areas V1, V2, V3, V4, V5, V4T, medial superior temporal area (MST), in lateral and inferior temporal areas TE1, TE2, TE3, TEO, TPO, superior temporal area (FST), PGa-IPa as well as in ventral temporal areas 35, 36, TF, entorhinal cortex, superior temporal rostral area (STR).

In addition, significant bilateral greater activations are also found in premotor areas 6Va and 6Vb and in motor area ProM of the left hemisphere as well as in lateral frontal areas 8Av, 45, 47 medial part (47M) and orbital part (47O), and more anterior in orbitofrontal cortex – area 11, area 13 medial (13M), area 13 lateral (13L), orbital periallocortex (OPAI) and orbital proisocortex (OPro). Finally, emotional faces elicit activations in a set of bilateral subcortical areas, including the superior colliculus (SC), periaqueductal gray (PAG), thalamus, medial geniculate nucleus (MGN), hippocampus, caudate, amygdala, substantia nigra (SNR) and in different parts of the cerebellum (posterior lobe of the cerebellum and each left and right side of the cerebellum) (Figure 2A).

Next, we identified brain areas recruited during the observation of neutral and negative facial expression conditions individually compared to their scrambled versions (i.e., neutral faces > scrambled neutral faces contrasts and negative faces > scrambled negative faces). Figures 2B and C show group activation maps representing brain regions with higher activations for neutral (B) or negative (C) facial expressions compared to their scrambled versions. Overall, each of these contrasts elicits a common network, similar to the one described above for all emotional faces, with stronger activations in occipital (V1, V2, V3, V4, V5, V4T), temporal (TE1, TE2, TE3, TEO, TF, FST, TPO, PGa-IPa, 35, 36, entorhinal cortex), premotor (6Va, 6Vb), frontal and orbitofrontal (8Av, 45, 47, 47M, 47O, 13M, 13L, OPAI, Opro) cortices as well as in some subcortical areas (SC, PAG, thalamus, MGN, hippocampus, caudate, amygdala and cerebellum) (Figures 2B and 2C). For neutral faces, preferential activations are also found in frontal area 13M and OPAI as well as in the piriform cortex (Pir) and in the putamen as compared to their scrambled version (Figure 2B), whereas for angry faces stronger activations are also observed in temporal area STR, in motor area ProM and in orbitofrontal area 11 as well as in the SNR (Figure 2C).

#### Negative facial expression enhances brain activity along the occipitotemporal cortical axis and subcortical areas compared to neutral facial expression

Next, we identified those brain areas that are more active during the observation of negative facial expressions as compared to neutral ones (i.e., negative faces > neutral faces contrast).

Figure 3 shows the regions more activated by negative faces compared to neutral faces. As observed in the previous section, the overall broad topologies of the circuitries for each facial expression are similar (Figure 2) but we find critical differences when contrasting the two conditions (Figure 3). This analysis reveals significant bilateral greater activations for negative faces along the occipitotemporal axis in occipital areas (visual areas V1, V2, V3, V4, V5), in the dorsomedial area V6, in inferior temporal areas (TE1, TE2, TE3, TEO, TF, TH), in superior temporal areas (V4T, MST, TPO, PGa-Ipa) and in the fundus of superior temporal sulcus FST. We also observe stronger subcortical activations in the bilateral hippocampus, the right geniculate nuclei, the right SC, the bilateral hypothalamus, bilateral amygdala, and also in the brainstem.

**Figure 3.**
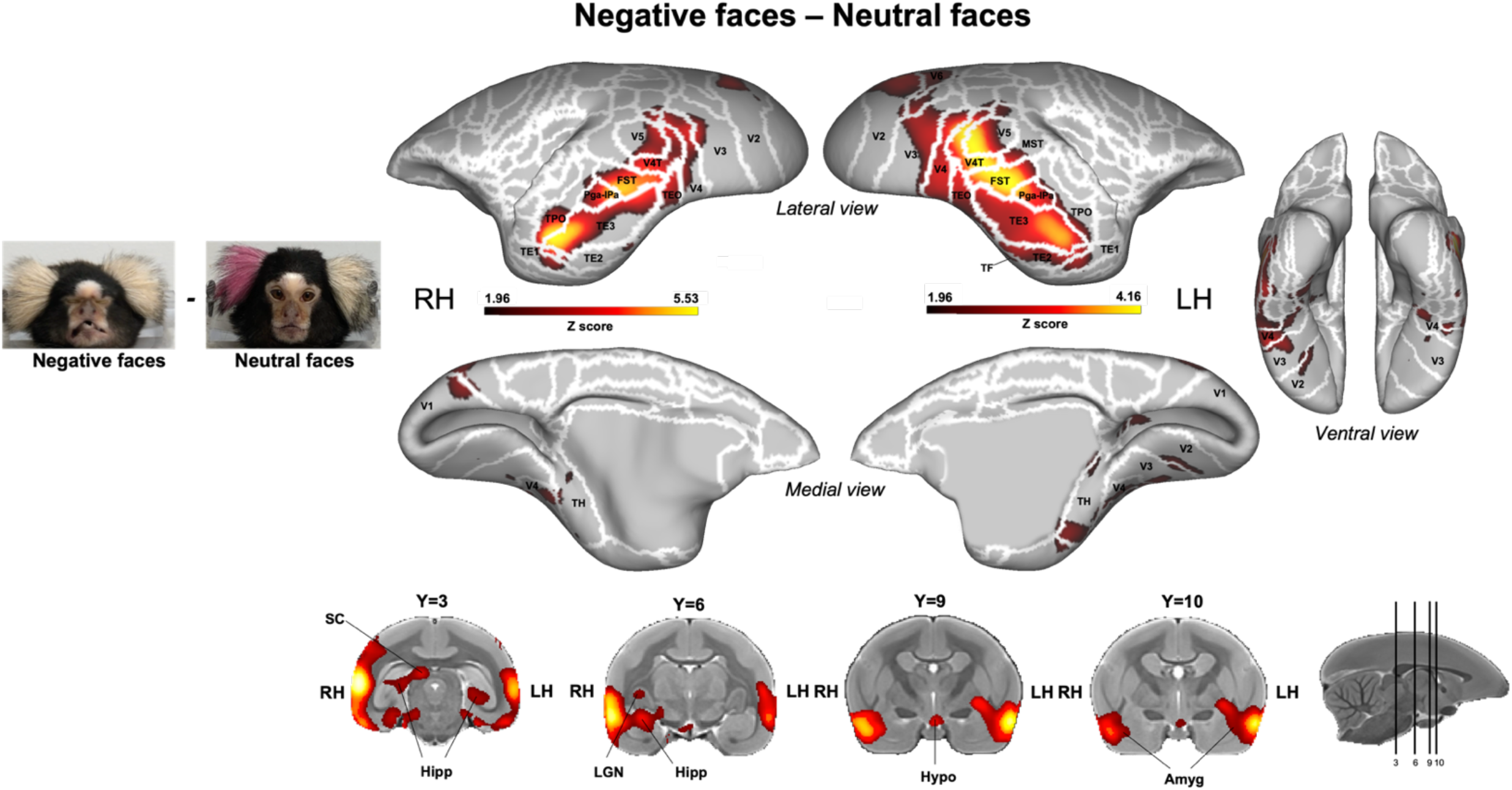
Brain network involved in negative facial expression processing compared to neutral facial expression. Group functional maps showing significant greater activations for negative compared to neutral facial expression displayed on the left and right fiducial marmoset cortical surfaces (lateral, medial and ventral views) as well as on coronal slices, to represent the activations in subcortical areas. The white line delineates the regions based on the Paxinos parcellation of the NIH marmoset brain atlas (Liu et al., 2018). The brain areas reported have activation threshold corresponding to z-scores > 1.96 (p<0.05, AFNI’s 3dttest++, cluster-size correction *α*=0.05 from 10000 Monte-Carlo simulations).

Overall, these results demonstrate that a set of brain areas is recruited for dynamic facial expressions processing but this network, in particular brain areas along the occipitotemporal axis and some subcortical areas like the amygdala, is stronger activated by negative compared to neutral facial expressions.

#### Negative and neutral facial expressions induce higher percentage signal change compared to their scrambled version in predefined face-patches ROIs

We selected temporal and frontal faces-patch areas previously defined in marmosets (Hung et al., 2015; Schaeffer et al., 2020): areas V2, V3, V4, TEO, V4T, FST, TE1, TE2, TE3, 45 and 47 (lateral, medial and orbital parts), which are also comparable to those found in humans and macaques for the processing of facial expressions (Fusar-Poli et al., 2009; Hadj-Bouziane et al., 2008). We also selected the amygdala known to be modulated by facial expressions in humans and macaques (Hadj-Bouziane et al., 2012; Vuilleumier and Pourtois, 2007). All regions were also present in our group activation maps (see above). Following the extraction of the time series from these ROIs, we compared the response magnitude between our conditions (Figure 4).

**Figure 4.**
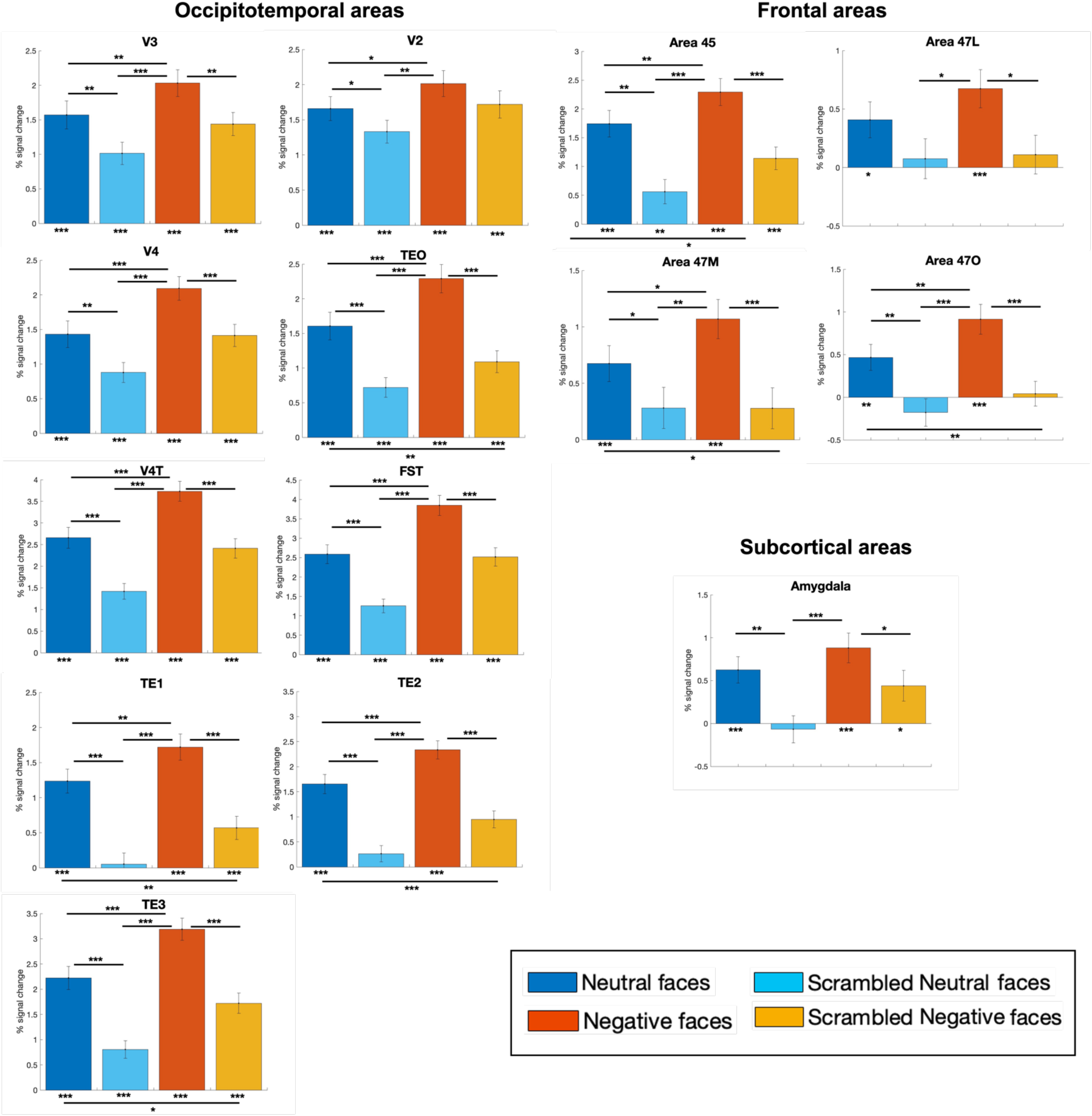
ROIs analysis: difference of percentage signal change responses between the four conditions (i.e., neutral and negative facial expressions and their scrambled versions). The magnitude of percentage signal change for each condition has been extracted from time series of 14 bilateral regions of interest using the Paxinos parcellation of the NIH atlas (Liu et al., 2018). The differences from baseline (represented by asterisks below each bar graph) and between conditions (represented by asterisk on horizontal bar) were tested using one sided paired t-tests corrected for multiple comparisons (FDR): p<0.05*, p<0.01** and p<0.001***. The error bars correspond to the standard error from the mean.

Most of these areas (12 ROIs) show significant stronger activations for neutral faces compared to scrambled neutral faces (paired t-tests with FDR correction, V3: t_(78)_=2.99 p=0.003; V2: t_(78)_=1.92 p=0.04; V4: t_(78)_=3.11 p=0.002; TEO: t_(78)_=4.30 p<0.001; V4T: t_(78)_=5.48 p<0.001; FST: t_(78)_=6.13 p<0.001; TE1: t_(78)_=5.14 p<0.001; TE2: t_(78)_=6.26 p<0.001; TE3: t_(78)_=6.19 p<0.001, Area 45: t_(78)_=4.43 p= 0.009; Area 47M: t_(78)_=1.93 p=0.05; Area 47O: t_(78)_=3.16 p=0.003, Amygdala: t_(78)_=3.14 p=0.02).

Almost the same areas also exhibit higher responses for negative faces compared to scrambled negative faces (paired t-tests with FDR correction, V3: t_(78)_=3.00 p=0.003; p<0.001; V4: t_(78)_=3.60 p<0.001; TEO: t_(78)_=5.67 p<0.001; V4T: t_(78)_=5.83 p<0.001; FST: t_(78)_=6.13 p<0.001; TE1: t_(78)_=5.67 p<0.001; TE2: t_(78)_=7.17 p<0.001; TE3: t_(78)_=7.02 p<0.001, Area 45: t_(78)_=4.42 p<0.001; Area 47L: t_(78)_=2.75 p=0.01; Area 47M: t_(78)_=3.82 p<0.001; Area 47O: t_(78)_=4.72 p<0.001, Amygdala: t_(78)_=2.26 p<0.001), except for V2 (t_(78)_=1.59 p=0.07).

All temporal and frontal regions (12 ROIs) are also significantly more activated by negative faces compared to neutral ones (paired t-tests with FDR correction, V3: t_(78)_=2.77 p=0.005; V2: t_(78)_=2.02 p=0.04; V4: t_(78)_=4.07 p<0.001; TEO: t_(78)_=3.85 p<0.001; V4T: t_(78)_=5.12 p<0.001; FST: t_(78)_=5.38 p<0.001; TE1: t_(78)_=2.61 p=0.007; TE2: t_(78)_=3.90 p<0.001; TE3: t_(78)_=4.92 p<0.001, Area 45: t_(78)_=2.51 p= 0.009; Area 47M: t_(78)_=2.15 p=0.03; Area 47O: t_(78)_=2.61 p=0.009). However, although sections of the amygdala were higher activated by negative compared to neutral faces in our group functional maps, it didn’t survive the FDR correction in the ROIs analysis (Amygdala: t_(78)_=1.25 p=0.1).

### Physiological results: increase of respiration rate with observation of negative facial expression

We recorded the monkeys’ respiration rate to determine if viewing negative facial expressions could influence their arousal state. In each run and for each animal, we computed the difference between the mean RR frequency obtained during the presentation of facial expression videos and the mean RR frequency during the baseline period (fixation dot) (i.e., delta RR, see methods). Figure 5 illustrates the animals’ delta RR frequency as a function of the viewing condition: neutral and negative facial expressions as well as scrambled version of each facial expression. The statistical analysis reveals a significant increase of delta RR for negative faces compared to scrambled negative faces (t_(16)_=1.93 p=0.04) and neutral ones (t_(16)_=1.85 p=0.04).

**Figure 5.**
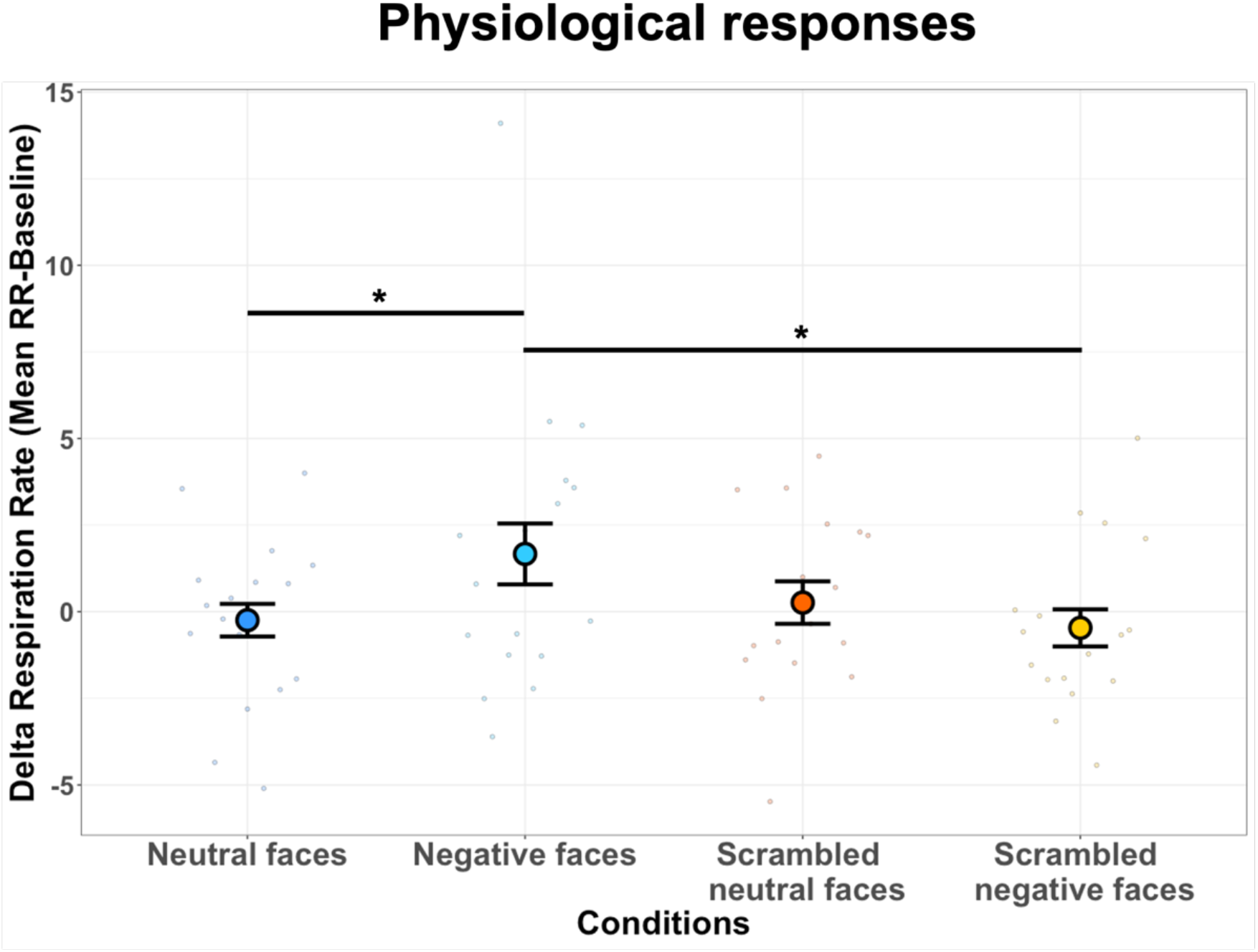
Respiration rate as a function of the viewing condition. Dot-plot depicting delta RR of all marmosets in bpm (i.e., mean RR during block of video clips – mean RR during baseline) according to each condition: neutral facial expression, negative facial expression, scrambled neutral faces and scrambled negative faces. In the plot, the mean delta RR is represented by the coloured dot (in blue for neutral faces, in light blue for negative faces, in red for scrambled neutral faces and in yellow for scrambled negative faces). The vertical bars represent the standard error from the mean and the small dots indicate individual values obtained for each run in all animals, for each condition. Three paired t-tests were performed: neutral faces *vs*. scrambled neutral faces, negative faces *vs*. scrambled negative faces and negative *vs*. neutral (p<0.05*, p<0.01** and p<0.001***)

As described in the method section, we only included the first run of each session in order to avoid habituation effect in physiological signals obtained by the repeated viewing of the negative facial expression across a same session, as already described in the literature for repeated exposures to fear stimuli (van Hout and Emmelkamp, 2002). Therefore, to control for this effect, we also performed the same analysis with only the last run of each session, and we computed the difference between each first vs. last runs conditions with paired t-tests. These results are presented in the supplementary materials (Figure S1). As predicted, the analysis reveals a significant decrease of delta RR for last runs compared to first runs only for the negative faces, suggesting a habituation effect with repeated exposure to negative faces in these animals. In order to identify if this effect could also impact neural responses, we also compared the response magnitude between first and last runs for each condition in the same ROIs previously selected (Figure S2 in supplementary materials). We observe that negative faces exhibit stronger activation for first compared to last runs in 7 ROIs (V3: t_(78)_=2.24 p=0.04; FST: t_(78)_=2.05 p=0.05; TE3: t_(78)_=2.73 p=0.03, Area 45: t_(78)_=3.26 p= 0.006; Area 47L: t_(78)_=2.94 p=0.02; Area 47O: t_(78)_=2.90 p=0.02), suggesting that our group maps may underreport the differences between negative and neutral faces.

## Discussion

In this study, we were interested in determining the brain network involved in the processing of dynamic facial expressions in New World marmosets. To this end, we presented videos of marmoset faces depicting either a neutral or negative facial expression and their scrambled versions during ultra-high field fMRI acquisitions. We also measured the respiration rate of the animals during the scans to determine if marmosets show physiological reactions to the videos. We observed that both neutral and negative dynamic face videos activated a large brain network including regions known to be implied in face processing. In addition, face videos activated motion-sensitive areas, premotor areas, and the posterior lobe of the cerebellum. Negative faces compared to neutral faces in particular enhanced brain activations in the temporal faces patches and in the amygdala, as well as in motion-sensitive areas, and were associated with higher respiration rates.

### Dynamic facial expressions processing recruit face-selective regions which are greater activated by negative facial expression

In humans, a number of fMRI studies have investigated the neural mechanisms underlying the processing of dynamic facial expressions (Arsalidou et al., 2011; Zinchenko et al., 2018). The occipitotemporal cortex including the fusiform gyrus, the anterior and posterior regions of the STS and the inferotemporal cortex as well as the dorsolateral prefrontal cortex, the orbitofrontal cortex and the amygdala are among the multiple structures that participate in recognizing facial emotions (Fusar-Poli et al., 2009; Haxby et al., 2000; Hennenlotter and Schroeder, 2006; Kanwisher et al., 1997; Posamentier and Abdi, 2003; Puce et al., 1998).

In Old World primates, these structures are situated along the occipitotemporal axis, mainly along the STS, in ventral and medial temporal lobe and in some subregions of the inferior temporal cortex, as well as in frontal cortex and some subcortical regions such as amygdala and hippocampus - regions known to be selective for faces processing (Bell et al., 2009; Hadj-Bouziane et al., 2008; Pinsk et al., 2005; Tsao et al., 2003; Weiner and Grill-Spector, 2015).

When we compare our activation pattern involved by dynamic facial expressions in marmosets, the regions along the occipitotemporal axis are comparable to those identified in macaques and humans (Tsao et al., 2008; Weiner and Grill-Spector, 2015). Our activations in posterior and anterior TE could correspond to the location of the STS in macaques and humans (Yovel and Freiwald, 2013). This suggests that, as in humans and macaque monkeys (Calder and Young, 2005), a division between dorsal and ventral axes appears for the processing of facial expressions in marmosets. Activations in V4T, FST, TE1, TE2, and TE3 may constitute a dorsal pathway which stands out compared to V4 and TEO areas which are more ventral areas. This suggest that the progression of facial expression-selective areas along this dorso-ventral axis may be preserved across Old World and New World primates.

Furthermore, our results are also similar to the temporal face-selective patches identified along the occipitotemporal axis in marmosets (Hung et al., 2015; Schaeffer et al., 2020): 1) V2/V3 (O patch), 2) V4/TEO (PV patch), 3) V4T/FST (PD patch), 4) posterior TE (MD patch) and 5) anterior TE (AD patch) as well as the frontal faces patches (Schaeffer et al., 2020). Moreover, our activations in orbitofrontal cortex (areas 11, 13M, 13L, OPAI and OPro) are in line with previous fMRI studies in humans (Adolphs, 2002; Beyer et al., 2015; Vuilleumier et al., 2001) and lesions studies in humans and macaques (Hornak et al., 1996; Willis et al., 2014) showing its direct role in recognition of facial expressions. This suggest that the frontal cortex in the marmoset brain could also participate to the perception and recognition of facial expressions.

Furthermore, our results also provide evidence of subcortical constituents (pulvinar, the hippocampus, the PAG, the SC, the LGN, and the amygdala) for facial expression processing in marmosets which is in line with humans and macaque studies demonstrating the modulation of amygdala by the viewing of facial expressions and emotional valence (Adolphs et al., 1994; Breiter et al., 1996; Brothers et al., 1990; Gothard et al., 2007; Hadj-Bouziane et al., 2012, 2008; Hoffman et al., 2007; Kuraoka and Nakamura, 2006; Vuilleumier and Pourtois, 2007). Also, it has been proposed that a subcortical pathway through the SC to the pulvinar could provide the amygdala with visual information about facial expressions allowing for a rapid and subconscious detection of emotions in Old World primates (Johnson, 2005; Tamietto and De Gelder, 2010). Therefore, we propose that a similar subcortical pathway involving the SC, the pulvinar and the amygdala could play an important role in the fast perception of emotional signals of faces in marmosets.

Finally, when we compare the activation patterns between negative and neutral faces, we find that visual and temporal regions are significantly more activated by negative faces, as previously reported in other species (Fusar-Poli et al., 2009; Hadj-Bouziane et al., 2008; Haxby et al., 2000; Hennenlotter and Schroeder, 2006; Hoffman et al., 2007; Kanwisher et al., 1997; Posamentier and Abdi, 2003; Puce et al., 1998). Moreover, the greater activations found in some subcortical areas, in particular in the amygdala, are in line with previous fMRI studies showing its preferential role in fearful and threatening face processing compared to other facial expressions (Hadj-Bouziane et al., 2008; Hoffman et al., 2007; Mattavelli et al., 2014; Morris et al., 1996). These findings suggest that negative facial expressions are processed similarly between species, implying specifically the occipitotemporal areas and the amygdala.

### Dynamic facial expressions processing involves regions outside face-selective regions

#### Motion-sensitive areas

In macaques, areas sensitive to “low-level” motion, such as the middle temporal area (MT/V5), area MST, and area FST participate in the representation of facial expression movements (Furl et al., 2012). In humans, static photographs with implied motion (Kourtzi and Kanwisher, 2000) or videos of facial expressions (Liang et al., 2017) also modulate responses in MT/V5 and MST. Similarly, our results show activations in areas MT, MST and FST for neutral and negative marmoset’s facial expressions compared to their scrambled versions. These regions were also more activate for negative faces as compared to neutral ones.

The role of motion-sensitive areas and face-selective areas is hypothesized to be distinct for dynamic facial expressions: motion-sensitive areas may represent expressions and face-selective areas may represent other facial attributes (Furl et al., 2015, 2012). Therefore, motion-sensitive areas activations could aid in the classification of facial expressions in marmosets.

#### Premotor areas

Interestingly, the contrasts between faces and scrambled faces also revealed activations in premotor area 6V. Electrical microstimulations in this area has been shown to evoke shoulder, back, neck, and facial movements in marmosets (Selvanayagam et al., 2019). In addition, this area has been shown to be activated by social interaction observation in marmosets (Cléry et al., 2021). Our results show that simply viewing facial expressions activates this area. This may be due to the facial mimicry phenomenon described as the process where facial expressions induce identical expressions in an observer (Nieuwburg et al., 2021). One of its consequences is the contagion of the emotional states of others which occur due to direct activation of the neural substrate involved in the experience of observed emotion. In human fMRI studies, greater activations have been identified in brain regions associated with facial movements (parts of primary motor and premotor cortex) and regions associated with emotional processing (insula and amygdala) during the perception of dynamic emotional facial expressions (Carr et al., 2003), which were also correlated with facial muscle responses (Rymarczyk et al., 2018).

Although our results seem to suggest that this contagion phenomenon may also apply to marmosets when simply viewing facial expression videos, it will be necessary to recording facial EMG responses to establish a role for area 6V in visual mimicry.

#### Cerebellum

In humans, the role of the cerebellum in motor functions has been well established. Recently, the number of studies associating the cerebellum with social cognition, in particular face and emotional processing, has increased substantially (Adamaszek et al., 2016). Recent studies in healthy individuals found that transcranial magnetic stimulation over the cerebellum affected participants’ abilities to accurately label emotional expressions (Ferrari et al., 2018) and significantly increased sensory processing in response to negative facial expressions (Ferrucci et al., 2012). In clinical populations, lesions studies have shown that cerebellar impairment was associated with deficits in emotional processing (Bolceková et al., 2017; Gold and Toomey, 2018). Meta-analysis of neuroimaging studies have also reported cerebellar activation in task involving emotional processing, in particular in the posterior lobe (vermal lobule VII), supporting an anterior sensorimotor *vs*. posterior emotional dichotomy in the human cerebellum (Baumann and Mattingley, 2012; Fusar-Poli et al., 2009; Sato et al., 2019; Stoodley and Schmahmann, 2009; Zinchenko et al., 2018). Therefore, our result of activation in the posterior part of the cerebellum during the observation of emotional faces in marmosets is in line with these human studies. However, we did not find the valence-specific cerebellar involvement during emotional faces processing as previously identified for negative faces in humans (Park et al., 2010; Schraa-Tam et al., 2012). Such differences were present when negative facial expressions were compared with positive facial expressions, which we did not include in our study. Future studies should explore a full range of facial expressions to determine if a dissociation of activation in cerebellum according to the valence of faces can also be observed in marmosets.

### Negative dynamic facial expression increases the level of arousal

Our results support a direct link between the observation of negative faces and an increase of RR in marmosets. This RR modulation was accompanied by higher responses in areas along the occipitotemporal axis, in some subcortical areas like the amygdala and in the brainstem. Interestingly, RR modulation was also linked to an increase of activation in the hypothalamus for negative faces – compared to neutral faces. It is well known that the hypothalamus has a central role in the integration of autonomic responses required for homeostasis and adaptation to internal or external stimuli allowing to monitor and regulate many variables in the body (Davis et al., 2009). Recently, an fMRI study in humans has demonstrated that exposure to a stress versus no-stress neutral experiment increased neural activation in brain circuits underlying the stress response (i.e., amygdala, hypothalamus, midbrain) which were accompanied by increased average heart rate and plasma cortisol response (Sinha et al., 2016). Therefore, physiological threats (e.g. emotional responses to negative faces) may be relayed to the occipitotemporal areas of the cerebral cortex and some subcortical areas, like the amygdala, via the stress-integrative brain centre located in the hypothalamus, influencing autonomic nervous system (ANS) activities in marmoset (Bains et al., 2015; McCorry, 2007). This activation of the sympathetic branch of the ANS would lead to an increased level of arousal, as observed in many previous physiological studies in humans (Bernat et al., 2006; Bradley et al., 2001; Fujimura et al., 2013; LANG et al., 1993).

### Conclusion

In summary, we report the first evidence from task-based fMRI for the existence of a cortico-subcortico-cerebellar network for dynamic facial expression processing in New World marmosets. Overall, our study reveals a network composed of faces-selective regions, motion-sensitive areas, premotor areas and the cerebellum that is involved in dynamic face processing and a set of temporal areas and the amygdala that are specifically recruited by the processing of negative faces which also increase RR. Therefore, we demonstrate that marmosets possess neural and physiological response to emotional faces, making them a powerful model for emotional face processing that is disrupted in neuropsychiatric and neurodevelopmental disorders (Azuma et al., 2015; Contreras-rodríguez et al., 2014; Dickstein and Xavier Castellanos, 2012; Harms et al., 2010; Monk et al., 2010; Safar et al., 2021; Uljarevic and Hamilton, 2013).

## Acknowledgements

Support was provided by the Canadian Institutes of Health Research (FRN 148365), the Canada First Research Excellence Fund to BrainsCAN, and a Discovery grant by the Natural Sciences and Engineering Research Council of Canada. We wish to thank Cheryl Vander Tuin, Whitney Froese, Hannah Pettypiece, and Miranda Bellyou for animal preparation and care, and Dr. Alex Li for scanning assistance.

## Notes

<u>Conflict of Interest</u> The authors declare that they have no conflict of interest.

### Competing Interest Statement

The authors have declared no competing interest.

## References

Adamaszek M, D’Agata F, Ferrucci R, Habas C, Keulen S, Kirkby KC, Leggio M, Mariën P, Molinari M, Moulton E, Orsi L, Van Overwalle F, Papadelis C, Priori A, Sacchetti B, Schutter DJ, Styliadis C, Verhoeven J. 2016. Consensus Paper: Cerebellum and Emotion. Cerebellum 2016 162 16:552–576. doi:10.1007/S12311-016-0815-8.

Adolphs R. 2002. Recognizing Emotion From Facial Expressions: Psychological and Neurological Mechanisms. Behav Cogn Neurosci Rev 1(1):21–62. doi:10.1177/1534582302001001003.

Adolphs R, Tranel D, Damasio H, Damasio A. 1994. Impaired recognition of emotion in facial expressions following bilateral damage to the human amygdala. Nat 1994 3726507 372:669–672. doi:10.1038/372669a0.

Aggleton JP, Passingham RE. 1981. Syndrome produced by lesions of the amygdala in monkeys (Macaca mulatta). J Comp Physiol Psychol 95:961–977. doi:10.1037/H0077848.

Albuquerque N, Guo K, Wilkinson A, Savalli C, Otta E, Mills D. 2016. Dogs recognize dog and human emotions. Biol Lett 12. doi:10.1098/RSBL.2015.0883.

Arsalidou M, Morris D, Taylor MJ. 2011. Converging evidence for the advantage of dynamic facial expressions. Brain Topogr 24:149–163. doi:10.1007/S10548-011-0171-4/FIGURES/5.

Azuma R, Deeley Q, Campbell LE, Daly EM, Giampietro V, Brammer MJ, Murphy KC, Murphy DG. 2015. An fMRI study of facial emotion processing in children and adolescents with 22q11.2 deletion syndrome. J Neurodev Disord 2015 71 7:1–16. doi:10.1186/1866-1955-7-1.

Bains JS, Cusulin JIW, Inoue W. 2015. Stress-related synaptic plasticity in the hypothalamus. Nat Rev Neurosci 2015 167 16:377–388. doi:10.1038/nrn3881.

Baumann O, Mattingley JB. 2012. Functional topography of primary emotion processing in the human cerebellum. Neuroimage 61:805–811. doi:10.1016/J.NEUROIMAGE.2012.03.044.

Bell AH, Hadj-Bouziane F, Frihauf JB, Tootell RBH, Ungerleider LG. 2009. Object representations in the temporal cortex of monkeys and humans as revealed by functional magnetic resonance imaging. J Neurophysiol 101:688–700. doi:10.1152/jn.90657.2008.

Bernat E, Patrick CJ, Benning SD, Tellegen A. 2006. Effects of picture content and intensity on affective physiological response. Psychophysiology 43:93. doi:10.1111/J.1469-8986.2006.00380.X.

Beyer F, Münte TF, Göttlich M, Krämer UM. 2015. Orbitofrontal Cortex Reactivity to Angry Facial Expression in a Social Interaction Correlates with Aggressive Behavior. Cereb Cortex 25:3057–3063. doi:10.1093/CERCOR/BHU101.

Blair RJR. 2003. Facial expressions, their communicatory functions and neuro-cognitive substrates. Philos Trans R Soc B Biol Sci 358:561. doi:10.1098/RSTB.2002.1220.

Bolceková E, Mojzeš M, Van Tran Q, Kukal J, Ostrý S, Kulišták P, Rusina R. 2017. Cognitive impairment in cerebellar lesions: A logit model based on neuropsychological testing. Cerebellum and Ataxias 4:1–11. doi:10.1186/S40673-017-0071-9/TABLES/6.

Bradley MM, Codispoti M, Cuthbert BN, Lang PJ. 2001. Emotion and motivation I: Defensive and appetitive reactions in picture processing. Emotion 1:276–298. doi:10.1037//1528-3542.1.3.276.

Breiter HC, Etcoff NL, Whalen PJ, Kennedy WA, Rauch SL, Buckner RL, Strauss MM, Hyman SE, Rosen BR. 1996. Response and Habituation of the Human Amygdala during Visual Processing of Facial Expression. Neuron 17:875–887. doi:10.1016/S0896-6273(00)80219-6.

Brothers L, Ring B, Kling A. 1990. Response of neurons in the macaque amygdala to complex social stimuli. Behav Brain Res 41:199–213. doi:10.1016/0166-4328(90)90108-Q.

Calder AJ, Young AW. 2005. Understanding the recognition of facial identity and facial expression. Nat Rev Neurosci 2005 68 6:641–651. doi:10.1038/nrn1724.

Carr L, Iacoboni M, Dubeaut MC, Mazziotta JC, Lenzi GL. 2003. Neural mechanisms of empathy in humans: A relay from neural systems for imitation to limbic areas. Proc Natl Acad Sci U S A 100:5497–5502. doi:10.1073/PNAS.0935845100/ASSET/386C6724-92B1-48DD-B65F-E3A5B48AD5BB/ASSETS/GRAPHIC/PQ0935845004.JPEG.

Ceccarini F, Caudek C. 2013. Anger superiority effect: The importance of dynamic emotional facial expressions. Vis cogn 21:498–540. doi:10.1080/13506285.2013.807901.

Cléry JC, Hori Y, Schaeffer DJ, Menon RS, Everling S. 2021. Neural network of social interaction observation in marmosets. Elife 10. doi:10.7554/ELIFE.65012.

Contreras-rodríguez O, Pujol J, Batalla I, Harrison BJ, Bosque J, Ibern-regàs I, Hernández-ribas R, Soriano-mas C, Deus J, López-solà M, Pifarré J, Menchón JM, Cardoner N. 2014. Disrupted neural processing of emotional faces in psychopathy. Soc Cogn Affect Neurosci 9:505. doi:10.1093/SCAN/NST014.

Cox RW. 1996. AFNI: Software for analysis and visualization of functional magnetic resonance neuroimages. Comput Biomed Res 29:162–173. doi:10.1006/cbmr.1996.0014.

Darwin C. 2004. The expression of the emotions in man and animals. Expr Emot man Anim. doi:10.1037/10001-000.

Davis JR, Giles AC, Rankin CH. 2009. Central Regulation of Autonomic Function. Encycl Neurosci 654–657. doi:10.1007/978-3-540-29678-2_904.

Dickstein DP, Xavier Castellanos F. 2012. Face processing in attention deficit/hyperactivity disorder. Curr Top Behav Neurosci 9:219–237. doi:10.1007/7854_2011_157/TABLES/1.

Ferrari C, Oldrati V, Gallucci M, Vecchi T, Cattaneo Z. 2018. The role of the cerebellum in explicit and incidental processing of facial emotional expressions: A study with transcranial magnetic stimulation. Neuroimage 169:256–264. doi:10.1016/J.NEUROIMAGE.2017.12.026.

Ferretti V, Papaleo F. 2019. Understanding others: Emotion recognition in humans and other animals. Genes, Brain Behav 18:e12544. doi:10.1111/GBB.12544.

Ferrucci R, Giannicola G, Rosa M, Fumagalli M, Boggio PS, Hallett M, Zago S, Priori A. 2012. Cerebellum and processing of negative facial emotions: cerebellar transcranial DC stimulation specifically enhances the emotional recognition of facial anger and sadness. Cogn Emot 26:786–799. doi:10.1080/02699931.2011.619520.

Fujimura T, Katahira K, Okanoya K. 2013. Contextual modulation of physiological and psychological responses triggered by emotional stimuli. Front Psychol 4:212. doi:10.3389/FPSYG.2013.00212/BIBTEX.

Furl N, Hadj-Bouziane F, Liu N, Averbeck BB, Ungerleider LG. 2012. Dynamic and Static Facial Expressions Decoded from Motion-Sensitive Areas in the Macaque Monkey. J Neurosci 32:15952–15962. doi:10.1523/JNEUROSCI.1992-12.2012.

Furl N, Henson RN, Friston KJ, Calder AJ. 2015. Network Interactions Explain Sensitivity to Dynamic Faces in the Superior Temporal Sulcus. Cereb Cortex 25:2876–2882. doi:10.1093/CERCOR/BHU083.

Fusar-Poli P, Placentino A, Carletti F, Landi P, Allen P, Surguladze S, Benedetti F, Abbamonte M, Gasparotti R, Barale F, Perez J, McGuire P, Politi P. 2009. Functional atlas of emotional faces processing: a voxel-based meta-analysis of 105 functional magnetic resonance imaging studies. J Psychiatry Neurosci 34:418.

Gold AK, Toomey R. 2018. The role of cerebellar impairment in emotion processing: A case study. Cerebellum and Ataxias 5:1–9. doi:10.1186/S40673-018-0090-1/TABLES/1.

Gothard KM, Battaglia FP, Erickson CA, Spitler KM, Amaral DG. 2007. Neural responses to facial expression and face identity in the monkey amygdala. J Neurophysiol 97:1671–1683. doi:10.1152/JN.00714.2006/ASSET/IMAGES/LARGE/Z9K0020779350008.JPEG.

Hadj-Bouziane F, Bell AH, Knusten TA, Ungerleider LG, Tootell RBH. 2008. Perception of emotional expressions is independent of face selectivity in monkey inferior temporal cortex. Proc Natl Acad Sci U S A 105:5591–5596. doi:10.1073/PNAS.0800489105.

Hadj-Bouziane F, Liu N, Bell AH, Gothard KM, Luh WM, Tootell RBH, Murray EA, Ungerleider LG. 2012. Amygdala lesions disrupt modulation of functional MRI activity evoked by facial expression in the monkey inferior temporal cortex. Proc Natl Acad Sci U S A 109. doi:10.1073/PNAS.1218406109.

Harms MB, Martin A, Wallace GL. 2010. Facial Emotion Recognition in Autism Spectrum Disorders: A Review of Behavioral and Neuroimaging Studies. Neuropsychol Rev 2010 203 20:290–322. doi:10.1007/S11065-010-9138-6.

Haxby J V., Hoffman EA, Gobbini MI, Haxby J V., Hoffman EA, Gobbini MI, Haxby J V., Hoffman EA, Gobbini MI. 2000. The distributed human neural system for face perception. Trends Cogn Sci 4:223–233. doi:10.1016/S1364-6613(00)01482-0.

Hennenlotter A, Schroeder U. 2006. Partly dissociable neural substrates for recognizing basic emotions: a critical review. Prog Brain Res 156:443–456. doi:10.1016/S0079-6123(06)56024-8.

Hoffman KL, Gothard KM, Schmid MCC, Logothetis NK. 2007. Facial-Expression and Gaze-Selective Responses in the Monkey Amygdala. Curr Biol 17:766–772. doi:10.1016/J.CUB.2007.03.040.

Hornak J, Rolls ET, Wade D. 1996. Face and voice expression identification in patients with emotional and behavioural changes following ventral frontal lobe damage. Neuropsychologia 34:247–261. doi:10.1016/0028-3932(95)00106-9.

Hung CC, Yen CC, Ciuchta JL, Papoti D, Bock NA, Leopold DA, Silva AC. 2015. Functional Mapping of Face-Selective Regions in the Extrastriate Visual Cortex of the Marmoset. J Neurosci 35:1160–1172. doi:10.1523/JNEUROSCI.2659-14.2015.

Johnson MH. 2005. Subcortical face processing. Nat Rev Neurosci 2005 610 6:766–774. doi:10.1038/nrn1766.

Johnston KD, Barker K, Schaeffer L, Schaeffer D, Everling S. 2018. Methods for chair restraint and training of the common marmoset on oculomotor tasks. J Neurophysiol 119:1636–1646. doi:10.1152/JN.00866.2017/SUPPL_FILE/SUPPLEMENTAL.

Kanwisher N, McDermott J, Chun MM. 1997. The Fusiform Face Area: A Module in Human Extrastriate Cortex Specialized for Face Perception. J Neurosci 17:4302–4311. doi:10.1523/JNEUROSCI.17-11-04302.1997.

Kourtzi Z, Kanwisher N. 2000. Activation in Human MT/MST by Static Images with Implied Motion. J Cogn Neurosci 12:48–55. doi:10.1162/08989290051137594.

Kuraoka K, Nakamura K. 2006. Impacts of facial identity and type of emotion on responses of amygdala neurons. Neuroreport 17:9–12. doi:10.1097/01.WNR.0000194383.02999.C5.

Lang PJ, Greenwald MK, Bradley MM, Hamm AO. 1993. Looking at pictures: Affective, facial, visceral, and behavioral reactions. Psychophysiology 30:261–273. doi:10.1111/J.1469-8986.1993.TB03352.X.

Liang Y, Liu B, Xu J, Zhang G, Li X, Wang P, Wang B. 2017. Decoding facial expressions based on face-selective and motion-sensitive areas. Hum Brain Mapp 38:3113. doi:10.1002/HBM.23578.

Liu C, Ye FQ, Yen CCC, Newman JD, Glen D, Leopold DA, Silva AC. 2018. A digital 3D atlas of the marmoset brain based on multi-modal MRI. Neuroimage 169:106–116. doi:10.1016/J.NEUROIMAGE.2017.12.004.

Liu N, Hadj-Bouziane F, Moran R, Ungerleider LG, Ishai A. 2017. Facial Expressions Evoke Differential Neural Coupling in Macaques. Cereb Cortex (New York, NY) 27:1524. doi:10.1093/CERCOR/BHV345.

Marcus DS, Harwell J, Olsen T, Hodge M, Glasser MF, Prior F, Jenkinson M, Laumann T, Curtiss SW, Van Essen DC. 2011. Informatics and data mining tools and strategies for the human connectome project. Front Neuroinform 5:4. doi:10.3389/FNINF.2011.00004/BIBTEX.

Mattavelli G, Sormaz M, Flack T, Asghar AUR, Fan S, Frey J, Manssuer L, Usten D, Young AW, Andrews TJ. 2014. Neural responses to facial expressions support the role of the amygdala in processing threat. Soc Cogn Affect Neurosci 9:1684–1689. doi:10.1093/SCAN/NST162.

McCorry LK. 2007. Physiology of the Autonomic Nervous System. Am J Pharm Educ 71. doi:10.5688/AJ710478.

Monk CS, Weng SJ, Wiggins JL, Kurapati N, Louro HMC, Carrasco M, Maslowsky J, Risi S, Lord C. 2010. Neural circuitry of emotional face processing in autism spectrum disorders. J Psychiatry Neurosci 35:105. doi:10.1503/JPN.090085.

Morris JS, Frith CD, Perrett DI, Rowland D, Young AW, Calder AJ, Dolan RJ. 1996. A differential neural response in the human amygdala to fearful and happy facial expressions. Nat 1996 3836603 383:812–815. doi:10.1038/383812a0.

Murphy FC, Nimmo-Smith I, Lawrence AD. 2003. Functional neuroanatomy of emotions: a meta-analysis. Cogn Affect Behav Neurosci 3:207–233. doi:10.3758/CABN.3.3.207.

Nieuwburg EGI, Ploeger A, Kret ME. 2021. Emotion recognition in nonhuman primates: How experimental research can contribute to a better understanding of underlying mechanisms. Neurosci Biobehav Rev 123:24–47. doi:10.1016/J.NEUBIOREV.2020.11.029.

Park JY, Gu BM, Kang DH, Shin YW, Choi CH, Lee JM, Kwon JS. 2010. Integration of cross-modal emotional information in the human brain: An fMRI study. Cortex 46:161–169. doi:10.1016/J.CORTEX.2008.06.008.

Parr LA, Dove T, Hopkins WD. 1998. Why Faces May Be Special: Evidence of the Inversion Effect in Chimpanzees. J Cogn Neurosci 10:615–622. doi:10.1162/089892998563013.

Pinsk MA, DeSimone K, Moore T, Gross CG, Kastner S. 2005. Representations of faces and body parts in macaque temporal cortex: A functional MRI study. Proc Natl Acad Sci U S A 102:6996–7001. doi:10.1073/PNAS.0502605102/SUPPL_FILE/02605FIG6.JPG.

Posamentier MT, Abdi H. 2003. Processing Faces and Facial Expressions. Neuropsychol Rev 2003 133 13:113–143. doi:10.1023/A:1025519712569.

Puce A, Allison T, Bentin S, Gore JC, McCarthy G. 1998. Temporal Cortex Activation in Humans Viewing Eye and Mouth Movements. J Neurosci 18:2188–2199. doi:10.1523/JNEUROSCI.18-06-02188.1998.

Russell JA. 1997. Reading emotions from and into faces: Resurrecting a dimensional-contextual perspective. Psychol Facial Expr 295–320. doi:10.1017/CBO9780511659911.015.

Rymarczyk K, Zurawski L, Jankowiak-Siuda K, Szatkowska I. 2018. Neural correlates of facial mimicry: Simultaneous measurements of EMG and BOLD responses during perception of dynamic compared to static facial expressions. Front Psychol 9:52. doi:10.3389/FPSYG.2018.00052/BIBTEX.

Safar K, Vandewouw MM, Taylor MJ. 2021. Atypical development of emotional face processing networks in autism spectrum disorder from childhood through to adulthood. Dev Cogn Neurosci 51:101003. doi:10.1016/J.DCN.2021.101003.

Sato W, Kochiyama T, Uono S, Sawada R, Kubota Y, Yoshimura S, Toichi M. 2019. Widespread and lateralized social brain activity for processing dynamic facial expressions. Hum Brain Mapp 40:3753. doi:10.1002/HBM.24629.

Schaeffer DJ, Gilbert KM, Hori Y, Gati JS, Menon RS, Everling S. 2019. Integrated radiofrequency array and animal holder design for minimizing head motion during awake marmoset functional magnetic resonance imaging. Neuroimage 193:126–138. doi:10.1016/J.NEUROIMAGE.2019.03.023.

Schaeffer DJ, Selvanayagam J, Johnston KD, Menon RS, Freiwald WA, Everling S. 2020. Face selective patches in marmoset frontal cortex. Nat Commun 2020 111 11:1–8. doi:10.1038/s41467-020-18692-2.

Schraa-Tam CKL, Rietdijk WJR, Verbeke WJMI, Dietvorst RC, Van Den Berg WE, Bagozzi RP, De Zeeuw CI. 2012. fMRI activities in the emotional cerebellum: A preference for negative stimuli and goal-directed behavior. Cerebellum 11:233–245. doi:10.1007/S12311-011-0301-2/TABLES/9.

Selvanayagam J, Johnston KD, Schaeffer DJ, Hayrynen LK, Everling S. 2019. Functional Localization of the Frontal Eye Fields in the Common Marmoset Using Microstimulation. J Neurosci 39:9197. doi:10.1523/JNEUROSCI.1786-19.2019.

Sinha R, Lacadie CM, Todd Constable R, Seo D. 2016. Dynamic neural activity during stress signals resilient coping. Proc Natl Acad Sci U S A 113:8837–8842. doi:10.1073/PNAS.1600965113/SUPPL_FILE/PNAS.1600965113.SAPP.PDF.

Smith SM, Jenkinson M, Woolrich MW, Beckmann CF, Behrens TEJ, Johansen-Berg H, Bannister PR, De Luca M, Drobnjak I, Flitney DE, Niazy RK, Saunders J, Vickers J, Zhang Y, De Stefano N, Brady JM, Matthews PM. 2004. Advances in functional and structural MR image analysis and implementation as FSL. Neuroimage 23:S208–S219. doi:10.1016/J.NEUROIMAGE.2004.07.051.

Stoodley CJ, Schmahmann JD. 2009. Functional topography in the human cerebellum: A meta-analysis of neuroimaging studies. Neuroimage 44:489–501. doi:10.1016/J.NEUROIMAGE.2008.08.039.

Strauss MM, Makris N, Aharon I, Vangel MG, Goodman J, Kennedy DN, Gasic GP, Breiter HC. 2005. fMRI of sensitization to angry faces. Neuroimage 26:389–413. doi:10.1016/J.NEUROIMAGE.2005.01.053.

Tamietto M, De Gelder B. 2010. Neural bases of the non-conscious perception of emotional signals. Nat Rev Neurosci 2010 1110 11:697–709. doi:10.1038/nrn2889.

Tate AJ, Fischer H, Leigh AE, Kendrick KM. 2006. Behavioural and neurophysiological evidence for face identity and face emotion processing in animals. Philos Trans R Soc B Biol Sci 361:2155. doi:10.1098/RSTB.2006.1937.

Tsao DY, Freiwald WA, Knutsen TA, Mandeville JB, Tootell RBH. 2003. Faces and objects in macaque cerebral cortex. Nat Neurosci 2003 69 6:989–995. doi:10.1038/nn1111.

Tsao DY, Moeller S, Freiwald WA. 2008. Comparing face patch systems in macaques and humans. Proc Natl Acad Sci U S A 105:19514–19519. doi:10.1073/PNAS.0809662105/SUPPL_FILE/0809662105SI.PDF.

Uljarevic M, Hamilton A. 2013. Recognition of emotions in autism: A formal meta-analysis. J Autism Dev Disord 43:1517–1526. doi:10.1007/S10803-012-1695-5/FIGURES/3.

Van Hout WJPJ, Emmelkamp PMG. 2002. Exposure in Vivo Therapy. Encycl Psychother 761–768. doi:10.1016/B0-12-343010-0/00091-X.

Vuilleumier P, Armony JL, Driver J, Dolan RJ. 2001. Effects of Attention and Emotion on Face Processing in the Human Brain: An Event-Related fMRI Study. Neuron 30:829–841. doi:10.1016/S0896-6273(01)00328-2.

Vuilleumier P, Pourtois G. 2007. Distributed and interactive brain mechanisms during emotion face perception: Evidence from functional neuroimaging. Neuropsychologia 45:174–194. doi:10.1016/J.NEUROPSYCHOLOGIA.2006.06.003.

Weiner KS, Grill-Spector K. 2015. The evolution of face processing networks. Trends Cogn Sci 19:240. doi:10.1016/J.TICS.2015.03.010.

Willis ML, McGrillen K, Palermo R, Miller L. 2014. The nature of facial expression recognition deficits following orbitofrontal cortex damage. Neuropsychology 28:613–623. doi:10.1037/NEU0000059.

Yovel G, Freiwald WA. 2013. Face recognition systems in monkey and human: are they the same thing? F1000Prime Rep 5. doi:10.12703/P5-10.

Zinchenko O, Yaple ZA, Arsalidou M. 2018. Brain Responses to Dynamic Facial Expressions: A Normative Meta-Analysis. Front Hum Neurosci 12:227. doi:10.3389/FNHUM.2018.00227/BIBTEX.

