## Supplementary materials for "Perception of dynamic facial expressions activates a cortico-subcortico-cerebellar network in marmosets"

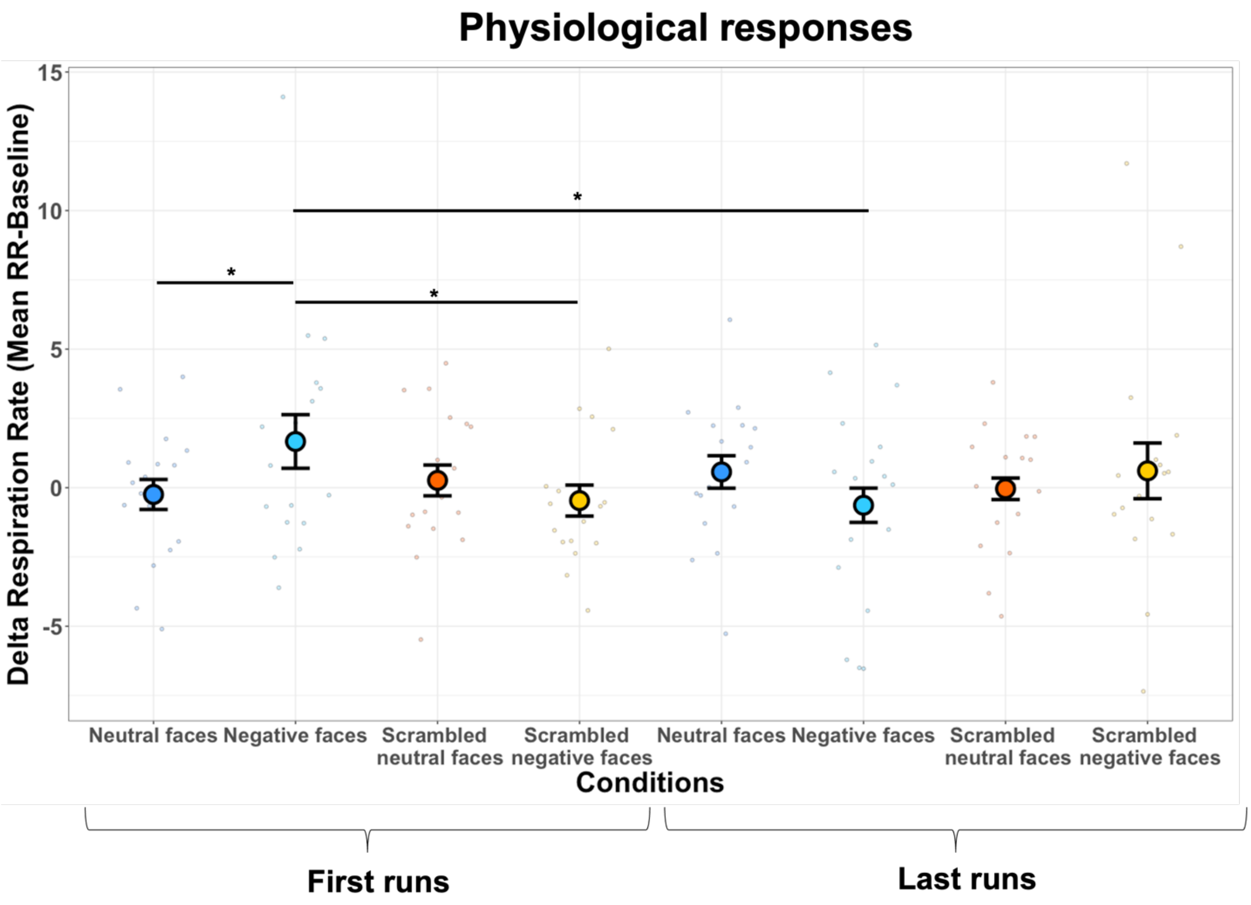


**Figure S1. Respiration rate as a function of the viewing condition for first *vs.* last runs of each session.** Dot-plot depicting delta Respiration Rate of all marmosets in bpm (RR, i.e., mean RR during block video clips conditions – mean RR during baseline) according to each condition: neutral facial expression, negative facial expression, scrambled neutral faces and scrambled negative faces for each first and last runs. In the plot, the mean delta RR is represented by the coloured dot (in blue for the neutral faces, in light blue for the negative faces, in red for the scrambled neutral faces and in yellow for the scrambled negative faces). The vertical bars represent the standard error from the mean and the small dots indicate individual values obtained for each run in all animals for each condition. Four paired t-tests were tested to identify difference between first and last runs conditions ($p<0.05$*, $p<0.01$** and $p<0.001$***).

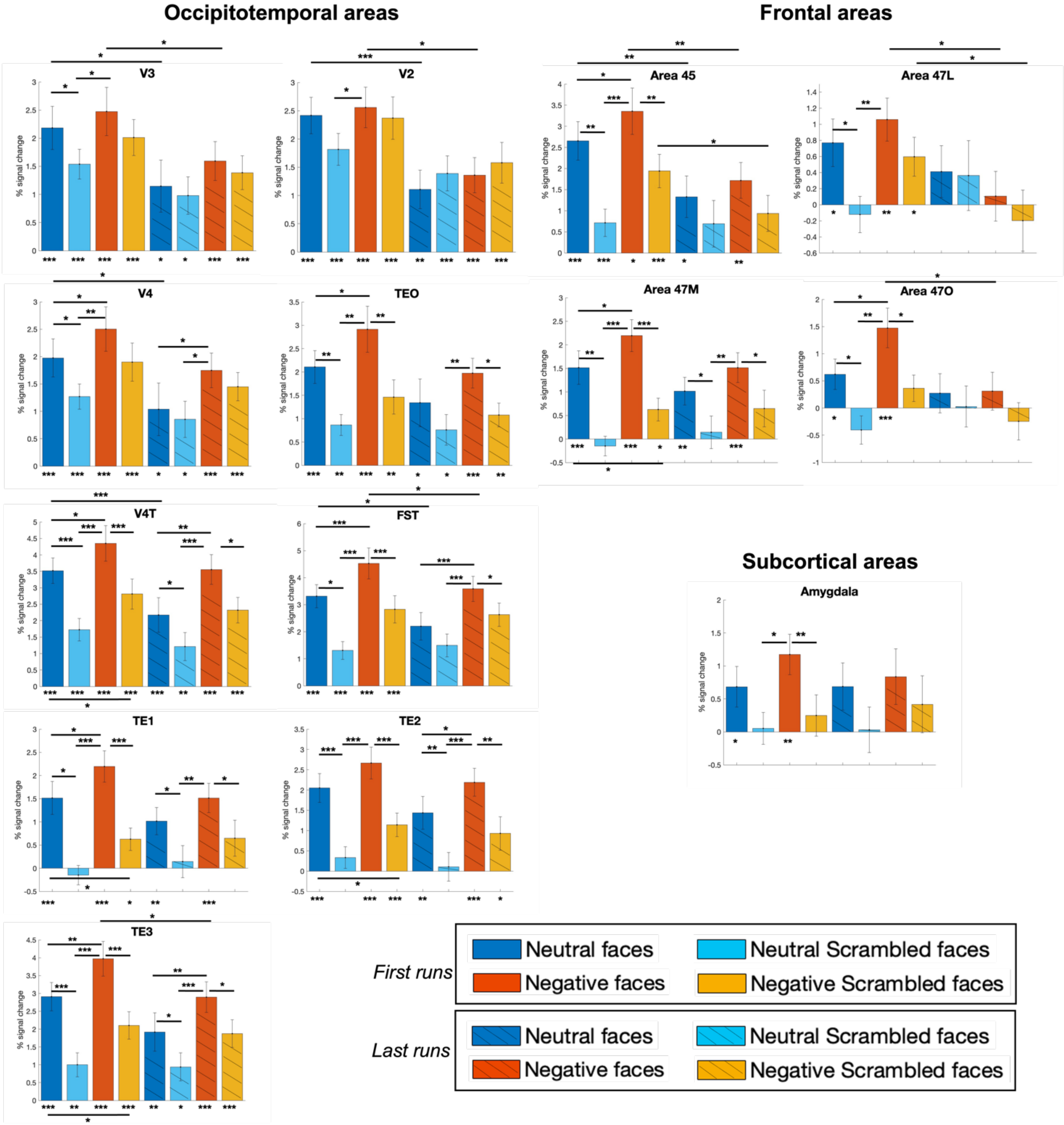
Occipitotemporal areas
Frontal areas
V3
V2
Area 45
Area 47L
V4
TEO
Area 47M
Area 47O
V4T
FST
Subcortical areas
Amygdala
TE1
TE2
TE3
% signal change
First runs
Neutral faces
Neutral Scrambled faces
Negative faces
Negative Scrambled faces
Last runs
Neutral faces
Neutral Scrambled faces
Negative faces
Negative Scrambled faces

**Figure S2. ROIs analysis comparing the magnitude of responses between conditions (i.e., neutral and negative facial expression and their scrambled versions) for first *vs.* last runs of each session.** The magnitude of percentage signal change for each condition has been extracted from time series of 14 bilateral regions of interest using the Paxinos parcellation of the NIH atlas (Liu et al., 2018). The differences from baseline (represented by asterisks below each bar graph) and between conditions inside first and last runs or between first and last runs (represented by asterisk on horizontal bar) were tested using one sided paired t-tests corrected for multiple comparisons (FDR): $p<0.05$*, $p<0.01$** and $p<0.001$***. Empty colored bar plots represent results from first runs whereas strikethrough colored bar plots represent results from last runs. The error bars correspond to the standard error from the mean.
